# Rapid Detection of *Burkholderia pseudomallei* with a Lateral Flow Recombinase Polymerase Amplification Assay

**DOI:** 10.1101/558262

**Authors:** Yao Peng, Zheng Xiao, Biao Kan, Wei Li, Wen Zhang, Jinxing Lu, Aiping Qin

**Affiliations:** Department of Pestis, National Institute for Communicable Disease Control and Prevention, Chinese Center for Disease Control and Prevention, Changping, Beijing, China; Department of Diarrheal Diseases, National Institute for Communicable Disease Control and Prevention, Chinese Center for Disease Control and Prevention, Changping, Beijing, China; Department of Bioinformatics, National Institute for Communicable Disease Control and Prevention, Chinese Center for Disease Control and Prevention, Changping, Beijing, China; Department of Hospital Antibiotics Resistance, National Institute for Communicable Disease Control and Prevention, Chinese Center for Disease Control and Prevention, Changping, Beijing, China; State Key Laboratory of infectious Diseases Prevention and Control, National Institute for Communicable Disease Control and Prevention, Chinese Center for Disease Control and Prevention, Changping, Beijing, China

## Abstract

Melioidosis is a severe infectious disease caused by gram-negative, facultative intracellular pathogen *Burkholderia pseudomallei* (*B. pseudomallei*). Although cases are increasing reported from other parts of the world, it is an illness of tropical and subtropical climates primarily found in southeast Asia and northern Australia. Because of a 40% mortality rate, this life-threatening disease poses a public health risk in endemic area. Early detection of *B. pseudomallei* infection benefits greatly to implement effective treatment timely, which is vital for prognosis of a melioidosis patient. In this study, a novel isothermal recombinase polymerase amplification combined with lateral flow dipstick (LF-RPA) assay was established for rapid detection of *B.pseudomallei*. A set of probe and primers targeting *orf*2 gene of *B. pseudomallei* were generated and parameters for the LF-RPA assay were optimized. Result can be easy visualized in 30 minutes with the limit of detection (LoD) as low as 20 femtogram (ca. 25.6 copies) of *B. pseudomallei* genomic DNA. The assay is highly specific as no cross amplification was observed with 35 non-*B. pseudomallei* pathogens. Isolates (*N*=19) from patients of Hainan province of China were retrospectively confirmed by the newly developed method. LoD for *B. pseudomallei* spiked soil and blood samples were 2.1×10^3^ CFU/g and 4.2×10^3^ CFU/ml respectively. Sensitivity of the LF-RPA assay was comparable to TaqMan Real-Time PCR, however, the LF-RPA assay exhibited a better tolerant to inhibitors in blood than the later. Our results showed that the LF-RPA assay is an alternative to existing PCR-based methods for detection of *B. pseudomallei* with a potentiality of early accurate diagnosis of melioidosis at point of care or in-field use.

## Introduction

Melioidosis, also called Whitmores disease, is an emerging infectious disease caused by the environmental bacterium *B. pseudomallei*. Although reported in many regions of the world, it is primary distributed in tropical and subtropical regions. Melioidosis is estimated to account for 89,000 deaths worldwide every year [1, 2]. *B. pseudomallei.* has been classified by the Centers for Disease Control and Prevention of US as a category B bioterrorism agent [3]. In China the epidemic areas of melioidosis are in Hainan, Guangdong and Guangxi province[4, 5]. Isolates from Hainan alone exhibited highly genetic diversity [6]. The main routes of *B. pseudomallei* infection are through inoculation of compromised skin, inhalation of contaminated soil during extreme weather event [7] and ingestion of contaminated water [8]. Manifestations of melioidosis are various and hard to differentiate from common pneumonia, flu or tuberculosis. Therefore, many cases have been under/misdiagnosed. In addition, *B. pseudomallei* is intrinsically resistance to a wide range of antibiotics, such as penicillin, ampicillin, first and second-generation of cephalosporins, gentamicin, tobramycin, streptomycin, and polymyxin[9]. A rapid and accurate diagnostic method is urgently needed.

Culture based method is the gold standard for diagnosing melioidosis, which typically requires 5-7 days in a highly equipped biosafety level 3 laboratory. This method has a limited diagnostic sensitivity, especially in the cases with prior antibiotic treatment. Molecular methods such as PCR and Real-Time PCR have been prevailed for diagnosis. However, they required sophisticated equipment as well as lengthy and complicated procedures. Finally, loop-mediated isothermal amplification (LAMP) assays were insensitive for detecting *B. pseudomallei* in blood samples [10].

Recombinase polymerase amplification (RPA) is a novel isothermal amplification method that can detect specific DNA or RNA with high sensitivity (less than 50 femtogram (fg) nucleic acid), short turnaround time (in 5-20 min) and few or no instrument needed. It has been utilized in detection of various pathogens including bacteria, viruses, parasites [11-15]. Basic RPA method consists of a primer pair which are able to scan and bind to the homology target DNA with the assistance of *E.coli Rec*A recombinase. The replication is achieved by DNA polymerase which owns a strand-displacement activity necessary to extend the oligonucleotides. The displaced DNA strand was stabilized by single-strand DNA binding proteins. DNA amplification can be conducted at ambient temperature and detected by agarose gel electrophoresis. Alternatively, when a third oligonucleotide probe is included, results can be analyzed by real time fluorescence or an oligochromotographic lateral flow strip. We described here the establishment of a TwishAmp™ nfo probe assay for detecting *orf*2 gene of *B. pseudomallei* and verified 19 clinical isolates of *B. pseudomallei*. The assay is quick and easy to perform with similar sensitivity to TaqMan Real-Time PCR (TaqMan PCR) but more tolerant to inhibitors in blood. The assay has exhibited its potential for detection of *B.pseudomallei* at point of need in endemic areas and emergence response in clinical settings.

## Materials and Methods

### Ethics Statement

Experimental protocols for collecting human clinical samples and isolating *B. pseudomallei* from the samples were approved by the Ethical Review Committee of the National Institute for Communicable Disease Control and Prevention (ICDC), Chinese Center for Disease control and Prevention (China CDC). Written consents have been obtained from all participants prior to the study.

### Primer and probe

The primers for basic RPA and probe for LF-RPA assay were designed to target *orf2*, one of type III secretion system cluster genes of *B. pseudomallei* (GenBank accession no. AF074878) following the instruction of TwistAmp® DNA amplification kits (TwistDx Ltd., UK). Primer pairs were initially screened against NCBI nucleotide database using BLAST, then evaluated via basic RPA. The probe used for the LF-RPA assay was a 46 bp length of nucleotides with FAM labeled at 5′ end, a tetrahydrofuran residue site (THF also referred as a dspacer) at 30 nucleotides downstream of the 5′ end and a block group (C3spacer) at 3′ end. Reverse primer for LP-RPA was conjugated with biotin at 5′ end. The primers for TaqMan PCR assay were selected as previously described [16]. All primers and probes were synthesized by Tianyihuiyuan.Co.Ltd (Beijing, China)(Table 1).

**Table 1.**
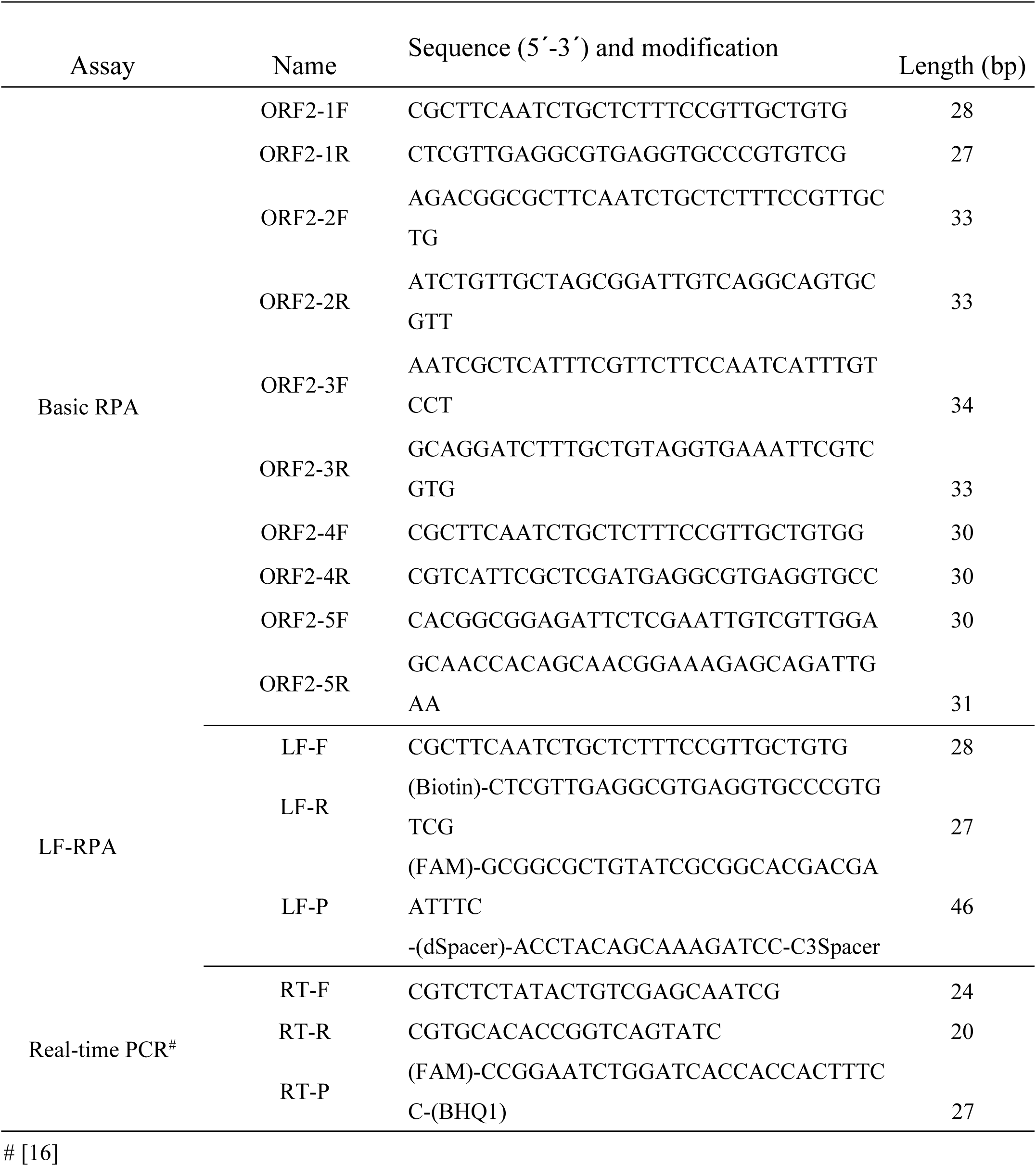
Sequences of primers and probes.

### Bacterial strains and genomic DNA preparation

All *B. pseudomallei* strains were handled in a China CDC certified Biosafety level 3 laboratory. DNA of standard *B. pseudomallei* strain (HN-Bp006) and 19 culture-confirmed *B. pseudomallei* strains isolated from blood sample of patients in Hainan province from 2016-2017, were extracted using the QIAamp DNA Mini Kit (Qiagen GmbH, Hilden, Germany). The DNA concentrations were quantified by the NanoDrop ND-1000 Spectrophotometers (Callibre, USA) and stored at −20°C until use. All *B. pseudomallei* strains were stocked at the Department of Yersinia pestis, National Institute for Communicable Disease Control and Prevention, Chinese Center for Disease Control and Prevention (China CDC).

### Basic RPA and later flow RPA (LF-RPA)

The basic RPA reaction was achieved by using the TwistAmp Basic kit (TwistDx, UK). A RPA reaction contained 2.4 μl forward primer (10 μM), 2.4 μl reverse primer (10 μM), 29.5 μl rehydration buffer, 12.2 μl H2O, 1 μl DNA. Amplification was initialized by adding of 2.5 μl Magnesium acetate (280 mM). The mixture was incubated at indicated temperature for 5 minutes, short vortex & spin, then returned to the water bath for an additional 15 minutes. The RPA product was purified by QIAquick PCR Purification kit (QIAGEN, Hilden, Germany) and analyzed on 1.5% agarose-gel. A series of basic RPA primers were tested.

LF-RPA assay entailed 2 primers and a FAM-labeled probe (Table 1) as described by TwistAmp nfo kit (TwistDX, Cambridge, UK). A LF-RPA reaction included 2.1 μl forward primer (10 μM), 2.1 μl reverse primer (10 μM), 29.5 μl rehydration buffer, 0.6 μl probe (10 μM), 12.2 μl H2O, 1 μl DNA, started by adding 2.5 μl of Magnesium acetate (280 mM). The reaction was run as previously described and stopped on ice. To detect amplicon, the reaction product was diluted at 1:50 in PBS. Result was visualized with a HybriDetect 2T dipstick following the manual (Milenia Biotec GmbH, Gießen, Germany).

### Temperature and time for the LF-RPA assay

The LF-RPA assays were performed with different reaction times to monitor the kinetic of amplification. Reactions were incubated for 5, 10, 15, 20, 25 and 30 minutes at 40 °C and stopped by placing reaction tubes on ice until further processing. Product was diluted in PBS and visualized as mentioned above. To find out the optimal amplification temperature, the LF-RPA assay was carried out with 2ng genomic DNA (gDNA) at different temperatures of 20°C, 25°C, 30°C, 37°C, 40°C, 45°C and 50 °C for 20 min. The experiment was repeated two times.

### Sensitivity and specificity of the LF-RPA assay

In order to assess the sensitivity of LF-RPA, gDNA of *B. pseudomallei* was 10-fold serial diluted. 1 μl of diluted DNA (2ng, 200 pg, 20 pg, 2pg, 200 fg, 20 fg, 2 fg) was used as template. The reactions were incubated at 40°C for 20 min. LoD was estimated as the lowest DNA amount, which is showing a clear band on the test strip. For comparison, the diluted DNA samples were tested in parallel by established TaqMan PCR protocol reported in a previous study[16].

To verify the specificity of the LF-RPA assay, gDNA(2ng) of the selected strains (*N*=5) and pools of 10 non-*B. pseudomallei* strains(*N*=30) (0.98-2.69ng/strain/reaction) were tested for possible cross reaction (Table 2). The experiment was carried out in triplicate and repeated twice.

**Table 2.** Non-*B. pseudomallei* bacterial strains used for specificity study.

Non- *B. pseudomallei* bacterial strains used for specificity study
| No. | Bacteria | Strains. | Concentration (ng/ $\mu$ l) | labeled on strip |
| --- | --- | --- | --- | --- |
| 1 | *Bacillus subtilis* | A186 | 18.6 | DNA-MIX1 |
| 2 | *Bacillus sturiens* | A170 | 17.7 |  |
| 3 | *Bacillus spore* | A1281 | 12.2 |  |
| 4 | *Shigella Bauer* | HBSH0002 | 11.7 |  |
| 5 | *Salmonella typhimurium* | JLSAL0002 | 12.5 |  |
| 6 | *Acinetobacter Bauman* | 301AB0145 | 11.7 |  |
| 7 | *Shigella flexneri* | HBSH001 | 14.9 |  |
| 8 | *Nissl spore* | A648 | 17.3 |  |
| 9 | *Staphylococcus aureus* | HBSA0007 | 17.4 |  |
| 10 | *Staphylococcus lugdunensis* | 3012SL0005 | 12.9 |  |
| 11 | *Salmonella bloom* | JLSA0003 | 16.2 | DNA-MIX2 |
| 12 | *Escherichia coli, EIEC* | HBEC0001 | 21.4 |  |
| 13 | *Bacillus sphaericus.* | A191 | 15.7 |  |
| 14 | *Bacillus licheniformis* | A180 | 11.1 |  |
| 15 | *Shigella Bauer* | JLSH0001 | 13.8 |  |
| 16 | *Shigella Song* | HBSH0006 | 10.9 |  |
| 17 | *Bacillus sphaericus.* | A184 | 17.0 |  |
| 18 | *Listeria monocytogenes* | ZJLM0051 | 18.0 |  |
| 19 | Drug resistant *Enterococcus* | NMVRE0001 | 11.3 |  |
| 20 | *Shigella dysentery* | HBSH0003 | 15.7 |  |
| 21 | *Bacillus licheniformis* | A197 | 12.1 |  |
| 22 | *Pantoea agglomerans* | ATCC27158 | 9.8 |  |
| 23 | *Bacillus cloacae* | A116 | 14.7 |  |
| 24 | *Kosotobacilli sakazakh* | ATCC29544 | 10.9 |  |
| 25 | *Pantoea spp* | A682 | 16.5 |  |
| 26 | *Pantoea ananas* | A729 | 26.9 |  |
| 27 | *Enterobacter sakazakii* | A278 | 12.7 |  |
| 28 | *Enterobacter aerogenes* | CGMCC1 | 18.6 |  |
| 29 | *Scattered fungi* | A895 | 13.0 |  |
| 30 | *Yersinia enterocolitica* | A398 | 11.0 |  |
| 31 | *Francisella tularensis* | 410062 | 2.0 | *F. tularensis* |
| 32 | *Francisella philomiragia* | A740 | 2.0 | *F. philomiragia* |
| 33 | *Burkholderia thailandensis* | W38 | 2.0 | *B. thailandensis* |
| 34 | *Yersinia pestis* | EV76 | 2.0 | *Y. pestis* |
| 35 | *Bacillus anthracis* | A714 | 2.0 | *B. ananthracis* |

### Detection of *B. pseudomallei* in spiked soil and blood sample by the LF-RPA

We collected soil samples (*N*=50) from different rice plantations in Zhanjiang Guangdong province in 2018. To isolate *B. pseudomallei* from the samples, 5-10g of the soil sample were mixed in 5-10 ml of Ashdown’s medium with gentamycin (4ug/ml final concentration) by vertexing, incubated at 40°C for 2 days and spun at 1,000rpm (939x g) for 5 minutes. Supernatant was centrifugated at 6,000rpm (3381x g) for 30 minutes then resuspended in 1ml of Ashdown medium. 100μl of the them were plated on a Ashdown plate (total of 3 plates) supplemented with gentamycin[17] then incubated at 40°C for 3-7 days. 600μl of the supernatant were centrifugated at 13,000rpm(15781x g) for 5min. Total DNA of the pellet was extracted by kit (Macherey-Nagel,GmßH&Co.KG,Germany) and concentration was measured (NanoDrop ND-1000). 1μl of the DNA (range from 3.7 to 25.6ng) was used for screening *B. pseudomallei* by TaqMan PCR. Selected samples were further verified by the LF-RPA assay.

Before spiking the soil or blood samples, *B. pseudomallei* colony forming units (CFU) was determined by plate-counting technique. Briefly, OD600=1 of *B. pseudomallei* culture was 10-fold serial diluted (10^-1^-10^-8^), aliquots of 20μl appropriate dilutions (10^-7^,10^-8^,10^-9^) were dripped onto LB plate in triplicate and incubated at 40°C for 48 hours for CFU counting. In order to estimate the LoD of the mocked soil samples, 100μl of each serial dilution (10^-3^-10^-8^) containing 4.2×10^5^ CFU, 4.2×10^4^ CFU, 4.2×10^3^ CFU, 4.2×10^2^ CFU 4.2×10^1^ CFU and 4.2 CFU of *B. pseudomallei* were mixed with 2g of soil sample in 900 μl of PBS. Total DNA of the spiked soil samples were extracted and tested by the LF-RPA and TaqMan PCR. LoD was expressed as the lowest CFU/g with which generates a visible band by the assay.

About one half of patients with melioidosis has positive blood cultures [18]. To determine the diagnostic potential of the LF-RPA assay, we spiked 100 μl of each serial diluted *B. pseudomallei* containing 4.2×10^5^ CFU, 4.2×10^4^ CFU, 4.2×10^3^ CFU, 4.2×10^2^ CFU 4.2×10^1^ CFU and 4.2 CFU of *B. pseudomallei* in 900 μl of defibrinated rabbit blood (Lefeikangtai,Beijing, China). DNA was extracted with Qiagen Blood &Tissue kit and used as template for the LF-RPA and TaqMan PCR. LoD was expressed as the lowest CFU/ml with which generates visible test line by the assay.

### Inhibition of the LF-RPA by blood

The inhibitory effect of substances in blood for the LF-RPA assay was investigated. Briefly, defibrinated rabbit blood or horse blood (Lefeikangtai,Beijing, China) were proportionally included in the LF-RPA and TaqMan PCR assay at the final concentration of 0%, 1%, 5% 10%, 12% and 15%. Standard LF-RPA and TaqMan PCR reactions were performed as mentioned above.

## Results

### Design and screening of primers

A 115-bp nucleotide sequence region of *orf*2, a gene within the type III secretion system gene cluster of the *B. pseudomallei* was previous targeted to distinguish *B. pseudomallei* from other microbial species by TaqMan PCR [16, 19]. A set of primers (Table1) were designed within the *orf*2 gene for basic RPA (GenBank accession no. AF074878) following the instruction of TwistAmp® DNA amplification kits (TwistDx Ltd., UK). In contrast to normal PCR, RPA primers require longer oligonucleotides, typically 30-35 base pair (bp) in order to stimulate homology recombination with the assistance of recombinase. Primers were paired and screened in a single tube reaction without the addition of the probe. Reaction conditions such as temperature, incubation time and primer concentration were optimized. As shown in Figure 1, ORF-1F/ORF-1R exhibited the best amplification efficiency (lane1) and therefore, was chosen in the follow-up experiments.

**Fig 1.**
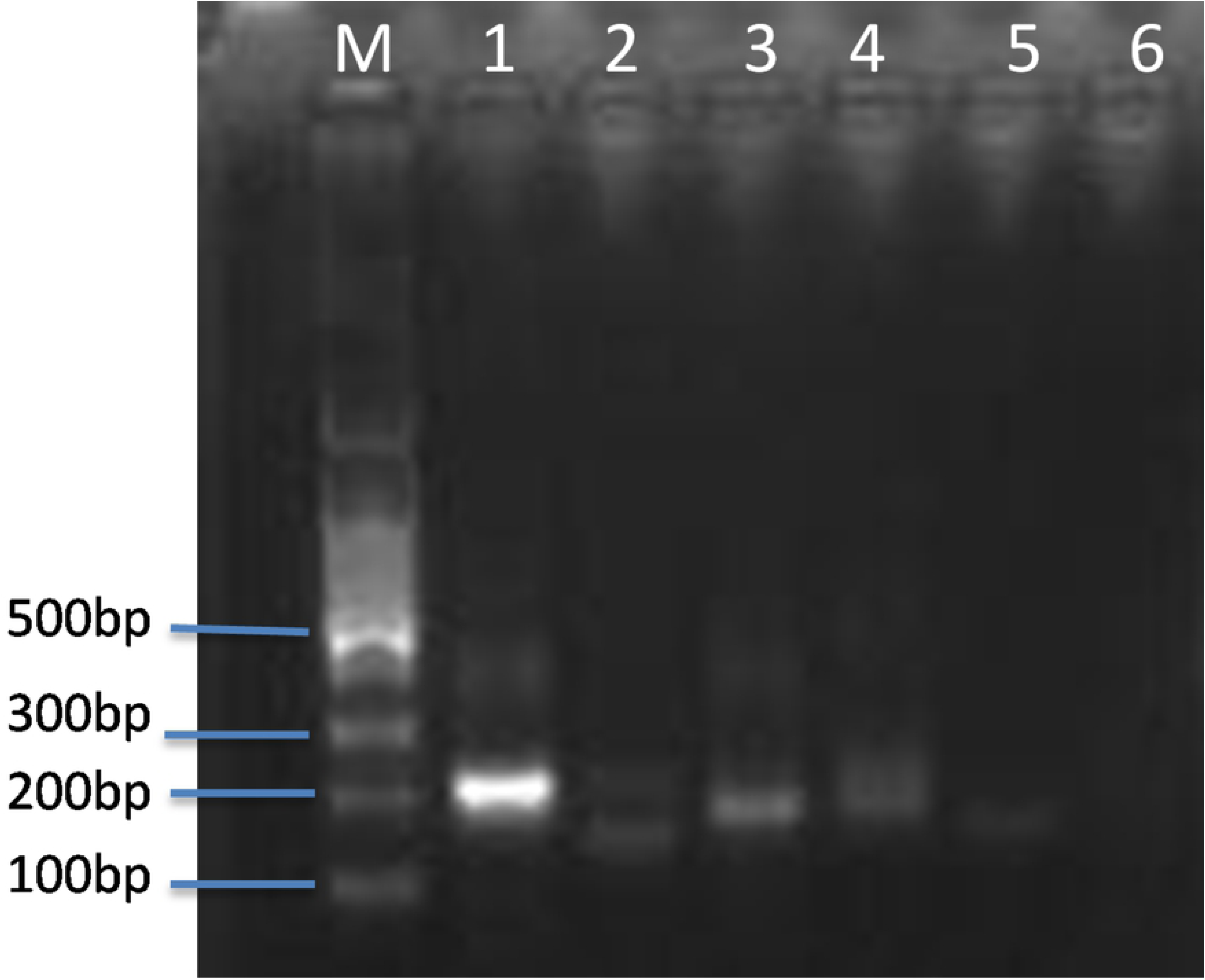
Primers screening for basic RPA. Several sets of primers targeting *orf2* gene of *B. pseudomallei* were tested for amplification efficiency by basic RPA (2ng gDNA as templated, at 40°C for 20 minutes). Lane 1-5 are primers of 1F/1R, 2F/2R, 3F/3R, 4F/4R, 5F/5R. lane 6 no primers control. M DNA ladder.

### Optimization of reaction temperature and time

To evaluate the optimal amplification temperature, LF-RPA assay was performed at indicated temperatures for 20 min (Figure 2A). The band density of test line on the strips varies with temperature over a wide range. The best result was achieved between at 30°C – 40°C. Therefore, the assay is suitable to body temperature, which is an advantage for operating the assay at low resource settings. The whole assay including RPA amplification and dipstick incubation takes 30 min or less.

**Figs 2A and 2B.**
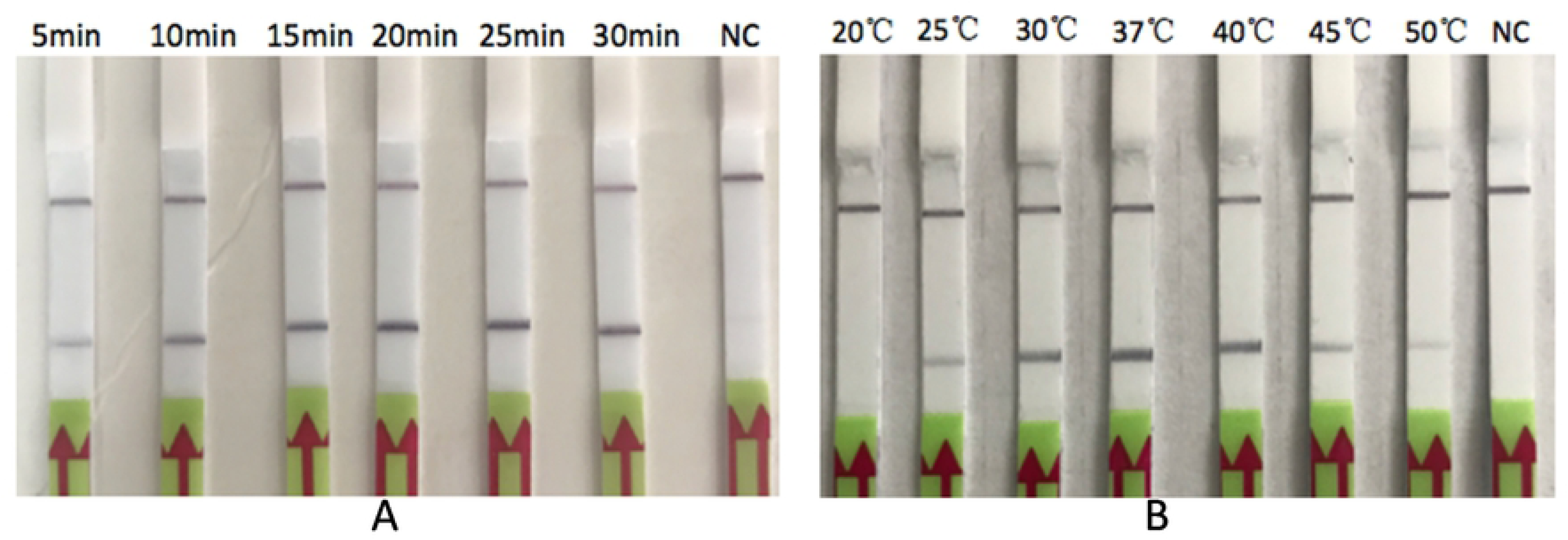
Optimization of temperature(A) and time(B) for the LF-RPA. gDNA of *B. pseudomallei. (*2ng) was used in each reaction at indicated temperatures for 20min (panel A) or for indicated time at 40°C (panel B). NC negative control. C, control line; T, test line. These experiments were repeated three times.

In order to monitor the kinetic of the LF-RPA amplification and estimate the optimal reaction time, the LF-RPA amplification was performed at 40°C for different incubation time. As the results show in Figure 2B, test band could be observed in as few as 5 min. The test line became intensified as time extended to 10 min or longer. Taking the detection efficiency and sensitivity into account, 15∼20 min amplification time works well for the LF-RPA assay.

### The specificity and sensitivity of the LF-RPA assay

To estimate the LoD of *B. pseudomallei.*, gDNA from 2 ng to 2 fg were tested by the LF-RPA assay. The results show our current method allow to detect as low as 20fg of gDNA (Figure 3A). Therefore, the sensitivity of LF-RPA is as good as the TaqMan PCR (Figure 3B).

**Figs 3A and 3B.**
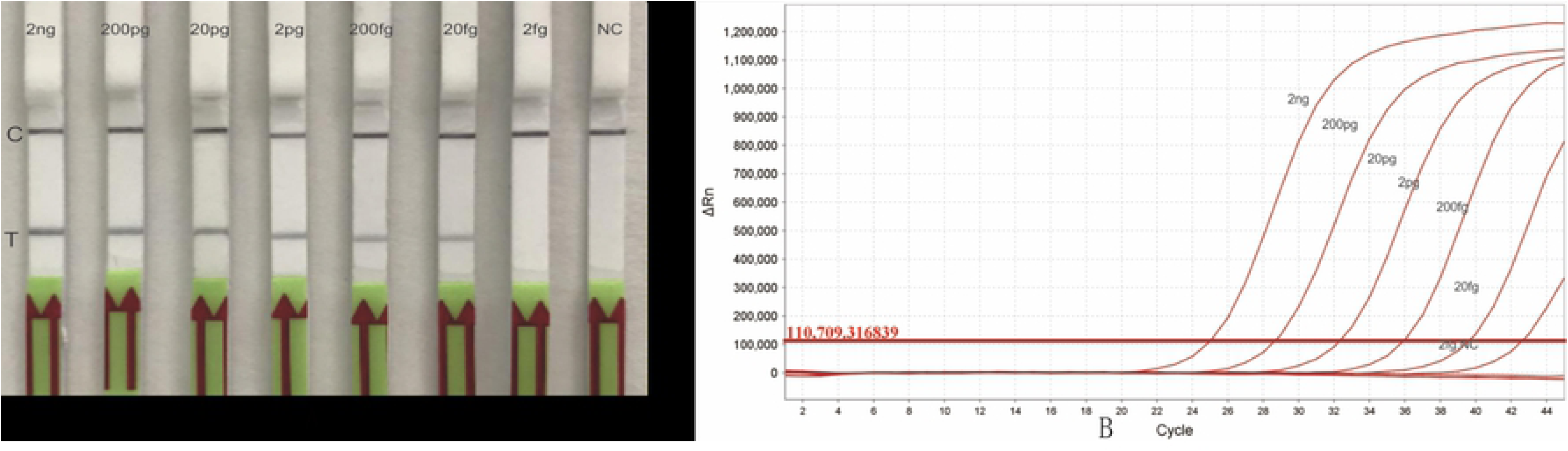
Sensitivity of the LF-RPA and Real-Time PCR. Serial diluted gDNA of *B. pseudomallei* (2ng to 2fg/reaction) was tested by LF-RPA at 40°C for 20 minutes (panel A) and by Real-Time PCR at 95°C for 5min, then 45 cycles of 95°C for 10 seconds, 60°C for 40 seconds (panel B). NC negative control C, control line; T, test line. This experiment was repeated twice with the same result.

To investigate the specificity of the LF-RPA, gDNA of a panel of bacterial pathogens were extracted individually and 2ng was used for the assay (Figure 4 lane 2, 3, 4, 5, 6). Positive result could be generated only with *B. pseudomallei* strain (lane 1), but not with the following bacteria: *B. thailandensis* (lane2), *Francisella tularensis* (lane 3), *Franciella philomirrihia* (lane 4), *Yersinia pestis* (lane 5) and *Bacillus anthracis* (lane 6). Pools of DNA from 10 non-*B. pseudomallei* pathogens were employed at final concentration of 0.98-2.69ng/each bacterium for possible cross reaction test (Table 2). As Figure 4 demonstrated the LF-RPA was able to correctly distinguished *B. pseudomallei* from other tested 35 pathogens, implying a high analytically specificity of this assay. Similar experiments were performed twice with the same results. Hence, the LF-RPA assay exhibited a trustworthy specificity.

**Fig 4.**
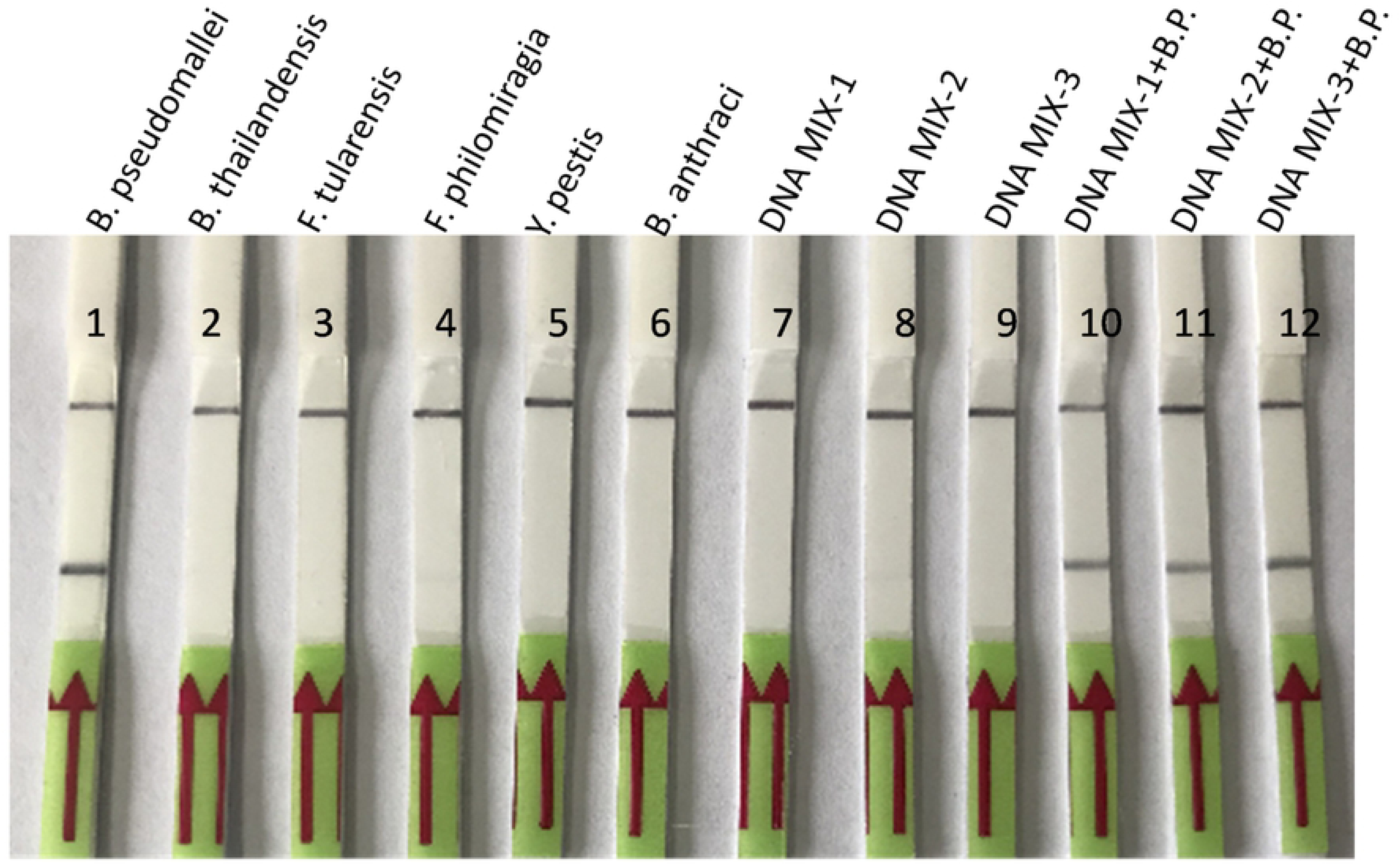
Specificity of the LF-RPA assay. LF-RPA reactions (at 40°C for 20 min.) were conducted with template of gDNA (2ng/reaction) of indicated bacteria (lane 1 to 6) or non-*B. pseudomallei* strains gDNA pools (0.98-2.69ng/each strain/reaction, lane 7, 8, 9). Lane 10, 11, 12 non-*B. pseudomallei* strains gDNA pools spiked with 2ng/reaction of HN-Bp006 gDNA respectively. Lane 1 positive control (HN-Bp006 gDNA). No cross-reaction was observed with 35 tested non-*B. pseudomallei* strains. C, control line; T, test line. This experiment was repeated three times with the same result.

### Validation of clinical isolates and evaluation of LF-RPA assay with *B. pseudomallei* spiked soil and blood samples

*B. pseudomallei* isolates (*N*=19) collected from clinical patients of Hainan province during 2016-2017 were formerly culture-confirmed. Here we further verified these strains by the LF-RPA assay. 100% of the isolates (19 /19) demonstrated a clearly visible band at the test line as the positive control (HN-Bp006 gDNA) indicating that the assay is a reliable method to detect *B. pseudomallei*. (Figure 5). This experiment was conducted two times with same result.

**Fig 5.**
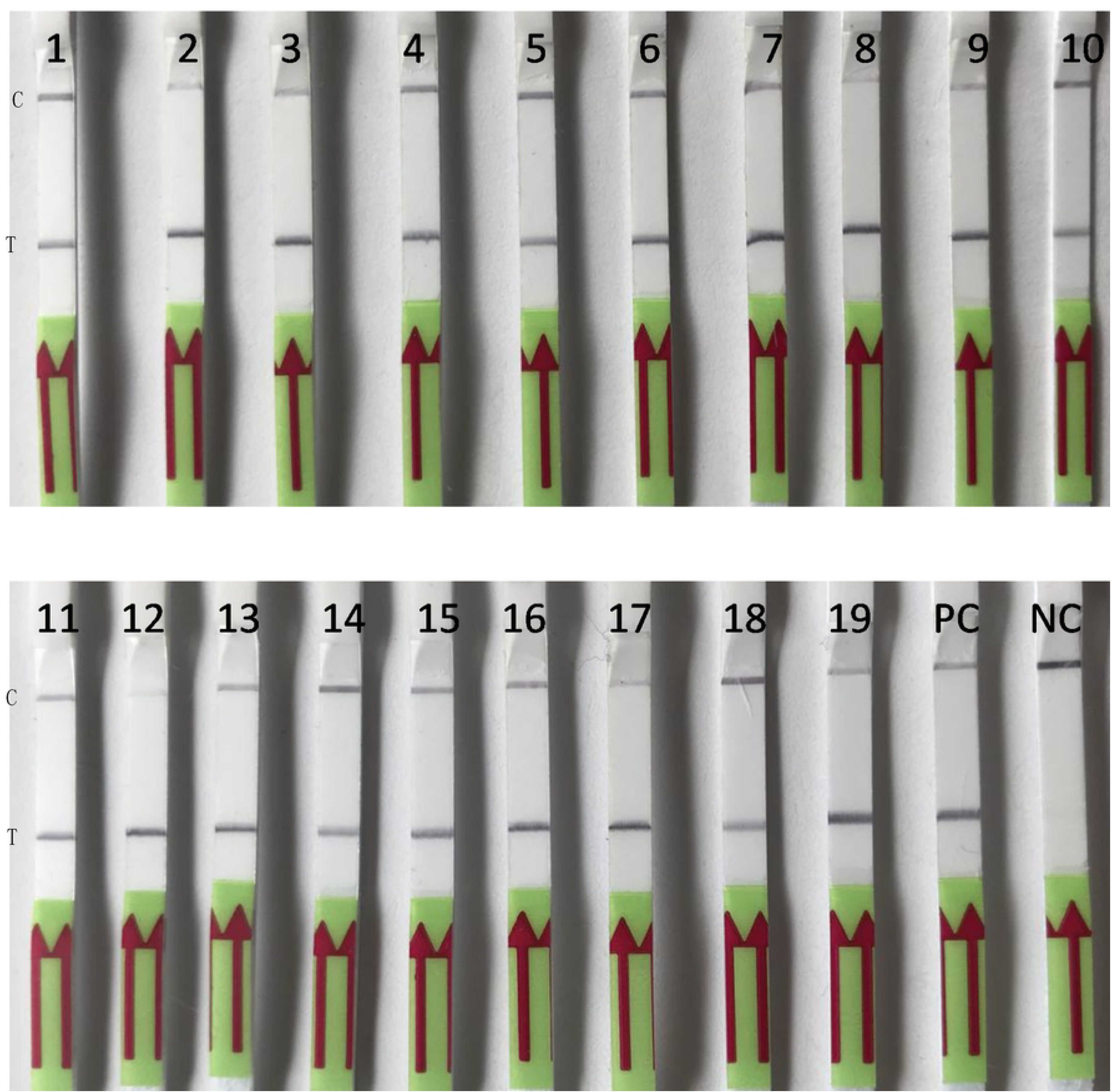
Analysis of clinical isolates by the LF-RPA assay. gDNA (2ng/reaction) of *B. pseudomallei* strains, collected from melioidosis patient from 2015 to 2017, were retrospectively confirmed by the LF-RPA (lane 1 to 19). PC positive control (2ng of HN-Bp006 gDNA), NC negative control (water). C, control line; T, test line.

*B. pseudomallei* is a soil dwelling pathogen. In order to evaluate the feasibility of the LF-RPA assay as a surveillance tool, 50 soil samples were collected from Guangdong province in 2018. DNA was prepared with soil DNA isolation kit as described in materials and methods. The LF-RPA assay and TaqMan PCR were conducted to detect *B. pseudomallei*. All the samples were TaqMan PCR negative (*N*=50). (S Figure 1). As expected, no *B. pseudomallei* was isolated from these samples by culture method. In this case, we spiked the soil samples with HN-Bp006 bacterial cells at different CFU/g to evaluate the capability of this assay to detect *B. pseudomallei* in soil sample. LoD of the LF-RPA and TaqMan PCR for spiked soil were both estimated at 2100 CFU/g (Figure 6).

**Fig 6.**
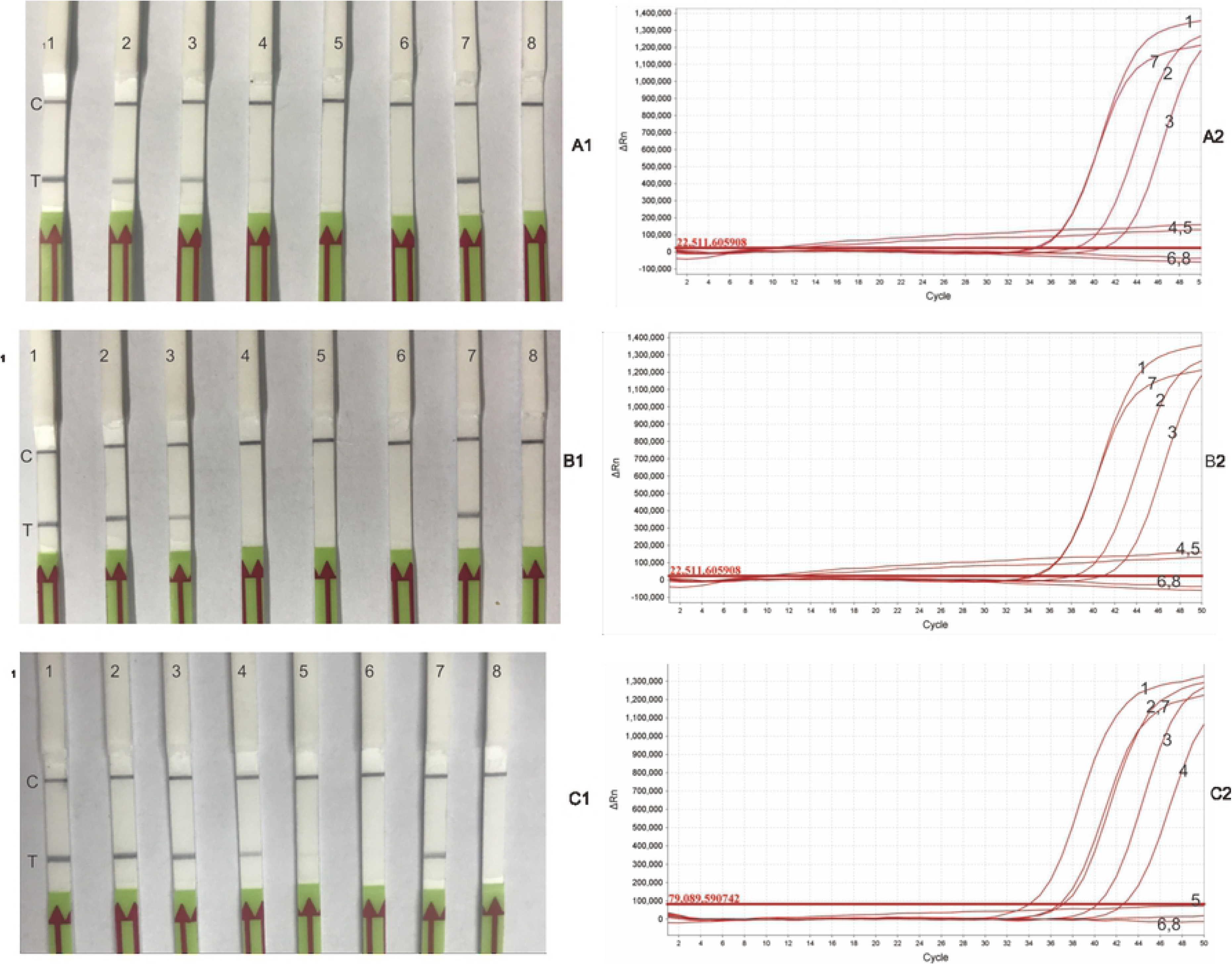
LoDs of the LF-RPA and Real-Time PCR. *B. pseudomallei* was 10-fold serial diluted and CFU/ml was estimated by plate-counting method. 100ul each of dilutions containing 4.2×10^5^ CFU, 4.2×10^4^ CFU, 4.2×10^3^ CFU, 4.2×10^2^ CFU 4.2×10^1^ CFU, and 4.2 CFU of *B. pseudomallei* were inoculated in 900ul of blood (A), 2g of soil (B) or 900ul of LB medium (C) respectively as described in materials and methods. gDNA was extracted by kits and used in a standard reaction of the LF-RPA (A1, B1, C1) and Real-Time PCR (A2, B2, C2). Lane1 4.2×10^5^, lane2 4.2×10^4^, lane3 4.2×10^3^, lane4 4.2×10^2^, lane5 4.2×10^1^, lane6 4.2. Lane7 2ng of HN-Bp006 gDNA, lane8 water control. LoD is the lowest CFU/ml or CFU/g that can be detected by the assays. C, control line; T, test line.

*B. pseudomallei* is commonly isolated from blood of melioidosis cases. To explore the possibility of the LF-RPA as a potential diagnostic means at point of care, *B. pseudomallei* was spiked in rabbit blood. The mocked clinical samples were tested by LF-RPA and TaqMan PCR (Figure 6). The LoD for both TaqMan PCR and the LF-RPA assay were 4.2×10^3^ CFU/ml, about 10-fold higher than the LoD of *B. pseudomallei* in culture medium (Table 3). We concluded that the capability of the LF-RPA assay to detect *B. pseudomallei* from mocked field samples is as effective as TaqMan PCR. Field samples such as blood or soil, may contain inhibitory substance(s) interfering both of LF-RPA and TaqMan PCR assay.

**Table 3.** LoDs of the LF-RPA and Real-Time PCR.

LoDs of the LF-RPA and Real-Time PCR
| Assay | LoD in |  |  |
| --- | --- | --- | --- |
|  | LB(CFU/ml) | Blood (CFU/ml) | Soil (CFU/g) |
| Real-Time PCR | 420 | 4200 | 2100 |
| LF-RPA | 420 | 4200 | 2100 |

### Effect of blood on the LF-RPA assay

The enzymatic nucleic acid amplification can be affected by numerous substances. To study the interference of blood on the LF-RPA assay, standard LF-RPA reactions (2ng template DNA, final volume of 50μl) were adjusted with rabbit or horse blood so that the percentage of blood within the reactions was 0%, 1%, 5%, 10%, 12% and 15% (v/v). The LF-RPA performed normally in the presence of blood below 12%. In contrast, TaqMan PCR lost detectable signal when percentage of blood in reaction volume exceeded 5% (Figure 7). Precipitation tends to form when percentage of blood is more than 1% in a standard PCR test (S. Figure 2). Increasing template in the assays (from100pg to 1ng) counteracts this inhibition in some extent (Table 4). Same inhibitory pattern was observed when horse blood was tested (data not shown), suggesting that this inhibition is not blood type dependent. In all, the LF-RPA assay tolerates the inhibitors presented in blood better than TaqMan PCR.

**Table 4.**
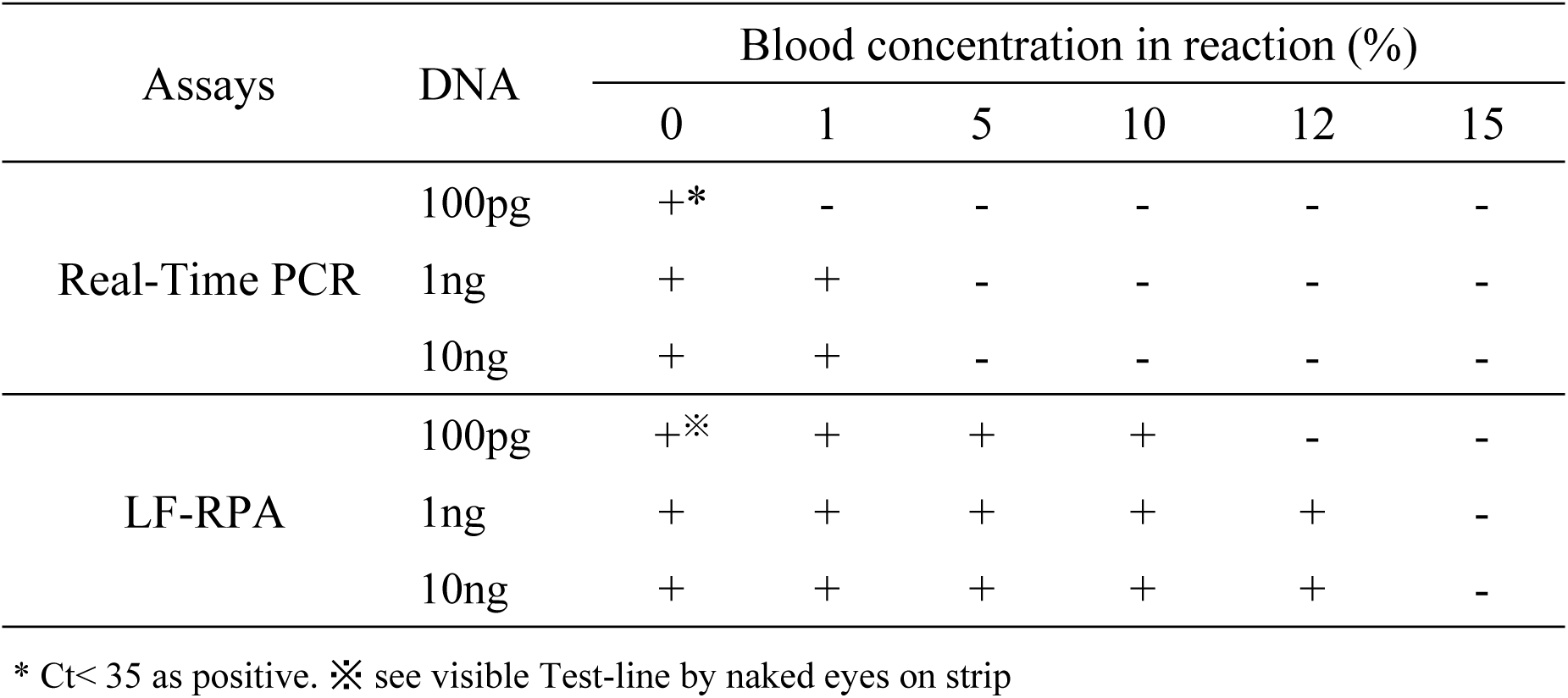
Inhibitory effect of blood on Real-Time PCR and the LF-RPA assay.

**Fig 7.**
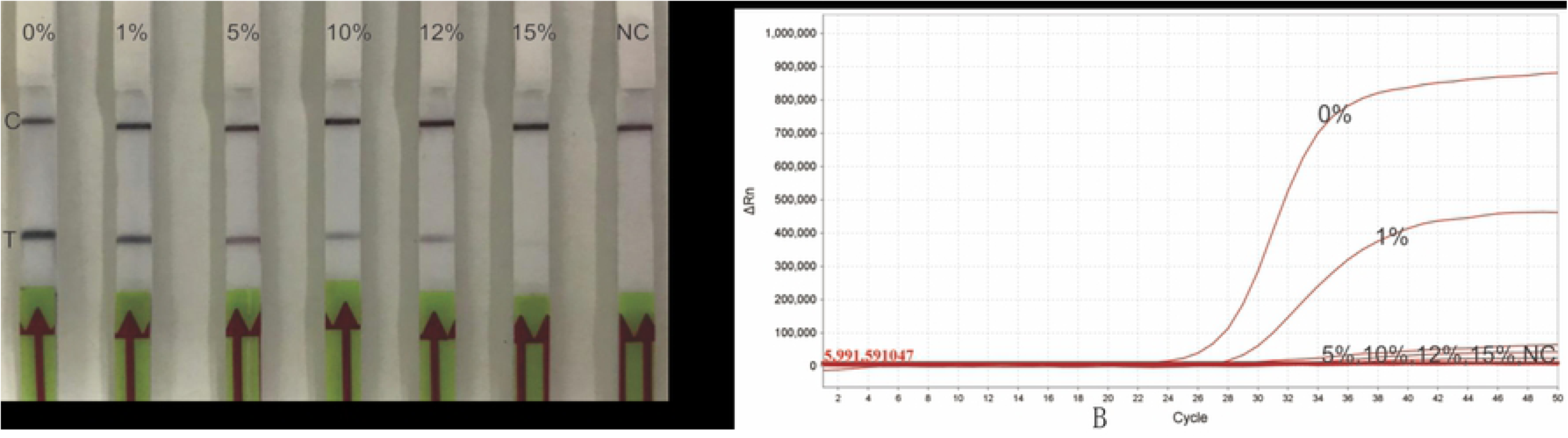
Effect of blood on the LF-RPA assay and Real-Time PCR. Defibrinated rabbit blood (or horse blood, data not shown) was included in reactions of the LF-RPA (panel A) or Real-Time PCR (panel B) at the indicated percentages. The LF-RPA was able to detect 1ng gDNA of *B. pseudomallei* at the presence of 12% of blood. Instead Real-Time PCR lost detectable signal when blood concentration in reaction was more than 1%. NC negative control. C, control line; T, test line.

## Discussion

First introduced in 2006, RPA represents an innovative DNA Isothermal detecting technology that has been used to detect a range of pathogens without the need of specific instruments. Amplification of target nucleic acid by this method can even be achieved under human body temperature [20], which is of a great advantage for field application. The LF-RPA assay developed here to detect *B. pseudomallei* is as sensitive as TaqMan PCR. This assay possesses a comparative LoD with LAMP [10]. The *orf2* gene was successfully illustrated in detection of *B. pseudomallei* of 19 clinical isolates but not of other 35 human pathogens, indicating the specificity of this gene for detecting of *B. pseudomallei*. In addition, the assay also can distinguish *B. pseudomallei* from the non-human pathogen of *B. thailandensis* [21]. The LF-RPA method is a very rapid technique, providing instructive information in less than 30 minutes (20 minutes reaction and 5minites detection) with a high sensitivity (a LoD of 25.6 copies). It is an alternative to existing PCR-based methods amenable to field-use.

The absence of typically clinical symptom of melioidosis renders early diagnosis difficult. Serology assays such as Indirect Hemagglutination Assay (IHA) remain the most widely used method for melioidosis diagnosis[22], but the sensitivity and specificity of those serological tests were not high enough for routine use[23]. Isolation of *B. pseudomallei* from patient is the conclusive diagnosis of melioidosis. It usually takes 5-7 days, in which time-frame patient may have been treated ineffectively and results in a higher fatality[24]. Immunofluorescence microscopy, PCR and TaqMan PCR methods were utilized in diagnosing clinical patient providing helpful evidence. However, they need expensive instruments and well-trained technical expertise that are not adapted to resource limited areas. RPA possesses an advantage over current molecular diagnosis methods, it may be able to detect clinical sample which had been missed by PCR[25]. Heamoglobin, lactoferrin and immunoglobulin G in blood for example are potential PCR inhibitors [26]. Residues of detergents, salts and ethanol et.al carried over during DNA isolation may hinder PCR reaction[27]. Although inhibited by whole blood[28], RPA had exhibited a certain tolerance to crude samples or crude materials with minimal processing [29]. The strategy reported here will increase diagnostic confidence and enable to make early accurate diagnosis of melioidosis.

Frequent specimens for *B. pseudomallei* include soil from environment and blood from clinical patient. The LF-RPA performed well with *B. pseudomallei* spiked soil and blood samples at a LoD of 2100cfu/g and 4200cfu/ml respectively. Although screened 50 soil specimens for *B. pseudomallei* collected from Guangdong province, known as a melioidosis endemic area in China, we foresee a need for further evaluation of the assay with large sample size and other clinical specimens such as pus, urine and body fluids. However, the satisfactory results of successful detection *B. pseudomallei* from mocked samples make us speculate that it is a promising method for clinical use.

A constraint of the LF-RPA for routine field application is the time-consuming DNA extraction steps involved in this study. However, not like TaqMan PCR, the LF-RPA demonstrated a better tolerance to the inhibitors existed in blood and other common PCR inhibitors [30]. We observed that when blood ratio was more than 1% (v/v) in a PCR or TaqMan PCR reaction, visible precipitation formed during the 95°C denature step. Also, the color of blood in reaction may interfere the result analysis of TaqMan PCR. On the other hand, LF-RPA is conducted at a constant lower temperature (40°C in this study) so bypassed this obstacle. Percentage of template in our routine LF-RPA reaction is 2% (1ul out of 50ul), therefore, blood sample is feasible to be applied as template directly.

False positive was reported for isothermal amplification methods[31]. Test band in the negative control was observed in this study. False positive is reduced when diluting the reaction products in PBS at ratio of 1 in 100 instead of 1 in 50 [13]. Dimer formation between probe and 3′ primer may play a role in this kind of assay noise [32]. In addition, we noticed that separate mixing of reagents and template areas greatly abrogates false positive. Lack of internal control impedes the practical application of this novel technique. This issue can be overcome by spiking external control during sample preparation [33] and developing of a duplex LF-RPA assay [34].

In summary, we first applied the isothermal recombinase polymerase amplification with later flow strip (LF-RPA) method for detection of *B. pseudomallei*. Although further studies are required to fully evaluated the utility of the assay in the field, it is a promising tool with advantages over currently available DNA diagnostic systems for melioidosis diagnosis.

## Acknowledgments

We would like to thank Dr. Shan Lu and Dr. Hongqing Zhao, Laboratory of Emergency Response, National Institute for Communicable Disease Control and Prevention, Chinese Center for Disease Control and Prevention, for their assistance with field samples collection. We thank Dr. Pierre Rivailler for reviewing this manuscript.

## Supporting information

**S1 Fig. Screening for *B. pseudomallei* from soil samples collected from Guangdong province.** Results of TaqMan PCR (50 samples, upper panel) or LF-RPA (selected 10 samples which ΔRn value elevated at Ct 20 in TaqMan PCR, lower panel). PC positive control (2ng of HN Bp-006 gDNA), NC negative control (water). C, control line; T, test line.

**S2 Fig. Formation of precipitation during PCR, but not LF-RPA when blood included in reaction.** Defibrinated rabbit blood was proportionally included in PCR reactions at the final concentration of 0%, 1%, 2% 3%, 4%, 5%,10% and 15%. PCR were performed at 95°C for 5min, then 30 cycles of 95°C for 10 seconds, 50°C for 30 seconds, 72°C for 30 seconds. The LF-RPA was conducted at 40 °C for 20 min. Precipitations were observed when percentage of blood was more than 1% in PCR reaction.

## References

1. Limmathurotsakul D, Golding N, Dance DAB, Messina JP, Pigott DM, Moyes CL, Rolim DB, Bertherat E, Day NPJ, Peacock SJ, Hay SI. Predicted global distribution of Burkholderia pseudomallei and burden of melioidosis. Nature Microbiology. 2016; 1(1). https://doi.org/10.1038/nmicrobiol.2015.8.

2. Dance DA, Limmathurotsakul D. Global Burden and Challenges of Melioidosis. Trop Med Infect Dis. 2018; 3(1). https://doi.org/10.3390/tropicalmed3010013 PMID: 30274411.

3. Lisa DR, Ali SK, Scott RL, Stephen MO, James MH. Public Health Assessment of Potential Biological Terrorism Agents. Emerging Infectious Disease journal. 2002; 8(2): 225. https://doi.org/10.3201/eid0802.010164.

4. Wang XM, Zheng X, Wu H, Zhou XJ, Kuang HH, Guo HL, Xu K, Li TJ, Liu LL, Li W. Multilocus Sequence Typing of Clinical Isolates of Burkholderia pseudomallei Collected in Hainan, a Tropical Island of Southern China. Am J Trop Med Hyg. 2016; 95(4): 760–764. https://doi.org/10.4269/ajtmh.16-0280 PMID: 27430537.

5. Dong S, Wu L, Long F, Wu Q, Liu X, Pei H, Xu K, Lu Y, Wang Y, Lin Y, Xia Q. The prevalence and distribution of Burkholderia pseudomallei in rice paddy within Hainan, China. Acta Tropica. 2018; 187: 165–168. https://doi.org/https://doi.org/10.1016/j.actatropica.2018.08.007.

6. Fang Y, Zhu P, Li Q, Chen H, Li Y, Ren C, Hu Y, Tan Z, Gu J, Mao X. Multilocus sequence typing of 102 Burkholderia pseudomallei strains isolated from China. Epidemiol Infect. 2016; 144(9): 1917–23. https://doi.org/10.1017/S0950268815003052 PMID: 26744829.

7. Cheng AC, Jacups SP, Gal D, Mayo M, Currie BJ. Extreme weather events and environmental contamination are associated with case-clusters of melioidosis in the Northern Territory of Australia. Int J Epidemiol. 2006; 35(2): 323–9. https://doi.org/10.1093/ije/dyi271 PMID: 16326823.

8. Limmathurotsakul D, Wongsuvan G, Aanensen D, Ngamwilai S, Saiprom N, Rongkard P, Thaipadungpanit J, Kanoksil M, Chantratita N, Day NP, Peacock SJ. Melioidosis caused by Burkholderia pseudomallei in drinking water, Thailand, 2012. Emerg Infect Dis. 2014; 20(2): 265–8. https://doi.org/10.3201/eid2002.121891 PMID: 24447771.

9. Wiersinga WJ, Currie BJ, Peacock SJ. Melioidosis. N Engl J Med. 2012; 367(11): 1035–44. https://doi.org/10.1056/NEJMra1204699 PMID: 22970946.

10. Chantratita N, Meumann E, Thanwisai A, Limmathurotsakul D, Wuthiekanun V, Wannapasni S, Tumapa S, Day NP, Peacock SJ. Loop-mediated isothermal amplification method targeting the TTS1 gene cluster for detection of Burkholderia pseudomallei and diagnosis of melioidosis. J Clin Microbiol. 2008; 46(2): 568–73. https://doi.org/10.1128/JCM.01817-07 PMID: 18039797.

11. Kim TH, Park J, Kim CJ, Cho YK. Fully integrated lab-on-a-disc for nucleic acid analysis of food-borne pathogens. Anal Chem. 2014; 86(8): 3841–8. https://doi.org/10.1021/ac403971h PMID: 24635032.

12. Liu L, Wang J, Zhang R, Lin M, Shi R, Han Q, Wang J, Yuan W. Visual and equipment-free reverse transcription recombinase polymerase amplification method for rapid detection of foot-and-mouth disease virus. BMC Vet Res. 2018; 14(1): 263. https://doi.org/10.1186/s12917-018-1594-x PMID: 30170587.

13. Rosser A, Rollinson D, Forrest M, Webster BL. Isothermal Recombinase Polymerase amplification (RPA) of Schistosoma haematobium DNA and oligochromatographic lateral flow detection. Parasit Vectors. 2015; 8: 446. https://doi.org/10.1186/s13071-015-1055-3 PMID: 26338510.

14. Vasileva Wand NI, Bonney LC, Watson RJ, Graham V, Hewson R. Point-of-care diagnostic assay for the detection of Zika virus using the recombinase polymerase amplification method. J Gen Virol. 2018; 99(8): 1012–1026. https://doi.org/10.1099/jgv.0.001083 PMID: 29897329.

15. Castellanos-Gonzalez A, White AC, Jr., Melby P, Travi B. Molecular diagnosis of protozoan parasites by Recombinase Polymerase Amplification. Acta Trop. 2018; 182: 4–11. https://doi.org/10.1016/j.actatropica.2018.02.002 PMID: 29452112.

16. Kaestli M, Richardson LJ, Colman RE, Tuanyok A, Price EP, Bowers JR, Mayo M, Kelley E, Seymour ML, Sarovich DS, Pearson T, Engelthaler DM, Wagner DM, Keim PS, Schupp JM, Currie BJ. Comparison of TaqMan PCR assays for detection of the melioidosis agent Burkholderia pseudomallei in clinical specimens. J Clin Microbiol. 2012; 50(6): 2059–62. https://doi.org/10.1128/JCM.06737-11 PMID: 22442327.

17. Howard K, Inglis TJJ. Novel Selective Medium for Isolation of Burkholderia pseudomallei. Journal of Clinical Microbiology. 2003; 41(7): 3312–3316. https://doi.org/10.1128/jcm.41.7.3312-3316.2003.

18. Limmathurotsakul D, Wuthiekanun V, Wongsuvan G, Pangmee S, Amornchai P, Teparrakkul P, Teerawattanasook N, Day NP, Peacock SJ. Repeat blood culture positive for B. pseudomallei indicates an increased risk of death from melioidosis. Am J Trop Med Hyg. 2011; 84(6): 858–61. https://doi.org/10.4269/ajtmh.2011.10-0618 PMID: 21633019.

19. Novak RT, Glass MB, Gee JE, Gal D, Mayo MJ, Currie BJ, Wilkins PP. Development and evaluation of a real-time PCR assay targeting the type III secretion system of Burkholderia pseudomallei. J Clin Microbiol. 2006; 44(1): 85–90. https://doi.org/10.1128/JCM.44.1.85-90.2006 PMID: 16390953.

20. Crannell ZA, Rohrman B, Richards-Kortum R. Equipment-free incubation of recombinase polymerase amplification reactions using body heat. PLoS One. 2014; 9(11): e112146. https://doi.org/10.1371/journal.pone.0112146 PMID: 25372030.

21. David ABD, Derek S, Erin PP, Direk L, Bart JC. Human Infection with <em>Burkholderia thailandensis</em>, China, 2013. Emerging Infectious Disease journal. 2018; 24(5): 953. https://doi.org/10.3201/eid2405.180238.

22. Sirisinha S, Anuntagool N, Dharakul T, Ekpo P, Wongratanacheewin S, Naigowit P, Petchclai B, Thamlikitkul V, Suputtamongkol Y. Recent developments in laboratory diagnosis of melioidosis. Acta Tropica. 2000; 74(2): 235–245. https://doi.org/https://doi.org/10.1016/S0001-706X(99)00076-5.

23. O’Brien M, Freeman K, Lum G, Cheng AC, Jacups SP, Currie BJ. Further Evaluation of a Rapid Diagnostic Test for Melioidosis in an Area of Endemicity. Journal of Clinical Microbiology. 2004; 42(5): 2239–2240. https://doi.org/10.1128/jcm.42.5.2239-2240.2004.

24. White NJ, Chaowagul W, Wuthiekanun V, Dance DAB, Wattanagoon Y, Pitakwatchara N. Halving of mortality of severe melioidosis by ceftazidime. The Lancet. 1989; 334(8665): 697–701. https://doi.org/https://doi.org/10.1016/S0140-6736(89)90768-X.

25. Bonney LC, Watson RJ, Afrough B, Mullojonova M, Dzhuraeva V, Tishkova F, Hewson R. A recombinase polymerase amplification assay for rapid detection of Crimean-Congo Haemorrhagic fever Virus infection. PLoS Negl Trop Dis. 2017; 11(10): e0006013. https://doi.org/10.1371/journal.pntd.0006013 PMID: 29028804.

26. Al-Soud WA, Radstrom P. Purification and characterization of PCR-inhibitory components in blood cells. J Clin Microbiol. 2001; 39(2): 485–93. https://doi.org/10.1128/JCM.39.2.485-493.2001 PMID: 11158094.

27. Santos SS, Nielsen TK, Hansen LH, Winding A. Comparison of three DNA extraction methods for recovery of soil protist DNA. Journal of Microbiological Methods. 2015; 115: 13–19. https://doi.org/https://doi.org/10.1016/j.mimet.2015.05.011.

28. Kersting S, Rausch V, Bier FF, von Nickisch-Rosenegk M. Rapid detection of Plasmodium falciparum with isothermal recombinase polymerase amplification and lateral flow analysis. Malaria Journal. 2014; 13(1): 99. https://doi.org/10.1186/1475-2875-13-99.

29. Silva G, Oyekanmi J, Nkere CK, Bomer M, Kumar PL, Seal SE. Rapid detection of potyviruses from crude plant extracts. Anal Biochem. 2018; 546: 17–22. https://doi.org/10.1016/j.ab.2018.01.019 PMID: 29378167.

30. Daher RK, Stewart G, Boissinot M, Bergeron MG. Recombinase Polymerase Amplification for Diagnostic Applications. Clin Chem. 2016; 62(7): 947–58. https://doi.org/10.1373/clinchem.2015.245829 PMID: 27160000.

31. Zasada AA, Zacharczuk K, Formińska K, Wiatrzyk A, Ziółkowski R, Malinowska E. Isothermal DNA amplification combined with lateral flow dipsticks for detection of biothreat agents. Analytical Biochemistry. 2018; 560: 60–66. https://doi.org/https://doi.org/10.1016/j.ab.2018.09.008.

32. Qi Y, Shao Y, Rao J, Shen W, Yin Q, Li X, Chen H, Li J, Zeng W, Zheng S, Liu S, Li Y. Development of a rapid and visual detection method for Rickettsia rickettsii combining recombinase polymerase assay with lateral flow test. PLoS One. 2018; 13(11): e0207811. https://doi.org/10.1371/journal.pone.0207811 PMID: 30475889.

33. Liu J, Platts-Mills JA, Juma J, Kabir F, Nkeze J, Okoi C, Operario DJ, Uddin J, Ahmed S, Alonso PL, Antonio M, Becker SM, Blackwelder WC, Breiman RF, Faruque ASG, Fields B, Gratz J, Haque R, Hossain A, Hossain MJ, Jarju S, Qamar F, Iqbal NT, Kwambana B, Mandomando I, McMurry TL, Ochieng C, Ochieng JB, Ochieng M, Onyango C, Panchalingam S, Kalam A, Aziz F, Qureshi S, Ramamurthy T, Roberts JH, Saha D, Sow SO, Stroup SE, Sur D, Tamboura B, Taniuchi M, Tennant SM, Toema D, Wu Y, Zaidi A, Nataro JP, Kotloff KL, Levine MM, Houpt ER. Use of quantitative molecular diagnostic methods to identify causes of diarrhoea in children: a reanalysis of the GEMS case-control study. The Lancet. 2016; 388(10051): 1291–1301. https://doi.org/10.1016/s0140-6736(16)31529-x.

34. Jauset-Rubio M, Tomaso H, El-Shahawi MS, Bashammakh AS, Al-Youbi AO, O’Sullivan CK. Duplex Lateral Flow Assay for the Simultaneous Detection of Yersinia pestis and Francisella tularensis. Anal Chem. 2018; 90(21): 12745–12751. https://doi.org/10.1021/acs.analchem.8b03105 PMID: 30296053.

